# Graphical pangenomics-enabled characterisation of structural variant impact on gene expression in Brassica napus

**DOI:** 10.1101/2024.03.14.585028

**Authors:** Agnieszka A. Golicz, Gözde Yildiz, Sven Weber, Tobias Kox, Amine Abbadi, Rod J. Snowdon, Silvia F. Zanini

## Abstract

Structural variants (SVs, eg. insertions and deletions) are genomic variations > 50 bp that are known to be associated with a range of crop traits, from yield to flowering behaviour and stress responses. Recently, pangenome graphs have emerged as a powerful framework for analysing genomic data by encoding population- or species-level diversity in one data structure. Pangenome graphs have the potential to serve as unbiased references for downstream applications, including SV genotyping and pan-transcriptomic analyses.

In this work, we hypothesized that extensive variation affects transcript quantification and expression quantitative trait locus (eQTL) analysis when relying on a single reference, and that using pangenome graphs can mitigate reference sequence bias.

We combined long and short read whole genome sequencing data with expression profiling of *Brassica napus* (oilseed rape) to assess the impact of SVs on gene expression regulation and explored the utility of pangenome graphs for eQTL analysis. We demonstrate that pangenome graphs provides a superior framework for eQTL analysis by eliminating single reference bias in gene expression quantification. Combined with the graph-based genotyping of SVs, we identified 240 eQTL-SVs found in close proximity of target loci. These SVs affect expression of genes related to important traits, are often not in linkage with SNPs and represent diversity unaccounted for in classical SNP-based analyses.

This study highlights the multiple advantages of graph-based approaches in population-scale studies and provides novel insight into gene expression regulation in an important crop.

## Introduction

Structural variants (SVs) are genomic alterations over 50 bp in length, with insertions and deletions representing the most common forms (Yildiz et al. 2023). SVs are prevalent in the complex genomes of major crops including wheat (Walkowiak et al. 2020), barley (Jayakodi et al. 2020) and oilseed rape (Chawla et al. 2021). Structural variants were also reported to be associated with important traits including yield and flowering time in oilseed rape (Song et al. 2020), fruit flavour in tomato (Li et al. 2023) and quality traits in cotton (Jin et al. 2023). SVs can alter gene function through several mechanisms including: changes to protein coding sequence, splicing patterns, gene expression levels, or any combination thereof (Zanini et al. 2022). One of the first large-scale studies reporting the wide-spread impact of structural variation on gene expression was Alonge et al. (2020), which used gene expression data from 23 accessions to link SVs to differential gene expression in tomato.

Expression quantitative trait loci (eQTL) analysis maps associations between genomic variation and gene expression. Results from eQTL studies are often used in conjunction with classical QTL mapping or genome wide association studies (GWAS) to pinpoint causal or candidate genes (Druka et al. 2010). They can however also be used to help understand the regulatory architecture of gene expression and complex phenotypic traits. The most common variants used in eQTL studies are single nucleotide polymorphisms (SNPs), however due to increasing capacity for population-scale SV discovery (Alonge et al. 2020; Chawla et al. 2021; Zhang et al. 2022), the impact of SVs on genome-wide expression patterns can now also be investigated in large scale eQTL analyses.

Recently, pangenome graphs have emerged as a powerful framework for the analysis of genomic data (Eizenga et al. 2020; Zanini et al. 2022). Pangenome graphs have the capacity to encode population-level genomic diversity in a single data structure, making it available as an unbiased reference for downstream applications. Graph-based genotyping of structural variants from short read data has become one of the most popular applications in crop plant research (Li et al. 2022; Liu et al. 2020). However, other important uses of pangenome graphs are emerging, including pan-transcriptomic analyses (Sibbesen et al. 2023; Coombes et al. 2024), where genomic variation is accounted for during mRNA-Seq read mapping and subsequent quantification.

Here, we hypothesized that the extensive genomic variation found in many crops can affect gene expression quantification and eQTL analysis when using a single reference genome, and that using a pangenome graph can help alleviate this problem.

In this study we combined long read Oxford Nanopore and short read Illumina genome sequencing data with mRNA-Seq data from young leaves of the major crop plant *Brassica napus* (oilseed rape) to assess the impact of structural variation on gene expression regulation and explore the utility of pangenome graphs for eQTL analysis (Fig 1A). We found that insertions, deletions and especially transposable elements (TEs) contribute to gene expression diversity and that functionally important SVs are frequently not in linkage disequilibrium with SNPs. We also found that using pangenome graphs as a reference can substantially improve transcript expression quantification.

**Figure 1.**
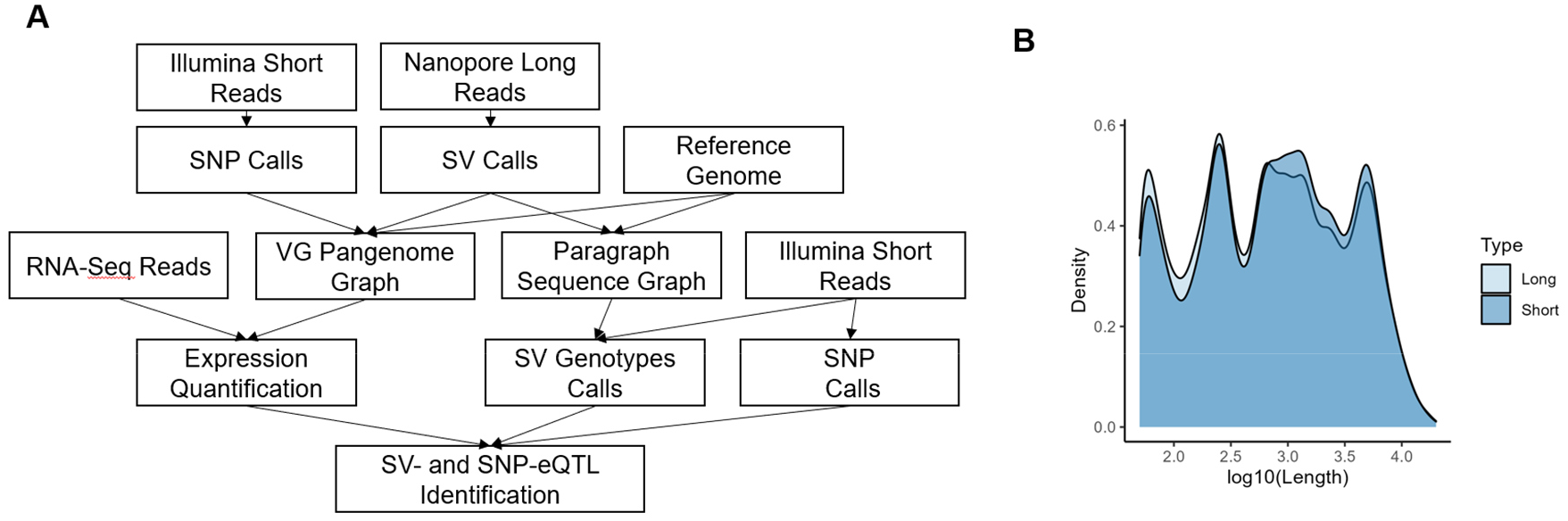
Overview of SV-eQTL analysis pipeline. **(A)** Long reads are used for SV identification. SVs along with SNPs identified from short reads are used for graph construction. Graphs are used for transcript expression quantification and SV genotyping using a larger collection of short read samples. **(B)** SVs identified from long reads and genotyped from short reads have similar length distribution.

## Results

### Long read sequencing of *Brassica napus* genotypes reveals tens of thousands of structural variants

We used Oxford Nanopore long read sequencing data for 57 lines and short read Illumina data for 100 lines (including the 57 lines selected for long read sequencing) of German winter oilseed rape (Fig 1A). The average coverage for long read data was 16.5x and for short read data 12x. The long read data was used to identify 48,396 insertions and 46,428 deletions. Short read data was used to genotype the SVs identified from long reads, using the graph-based genotyper Paragraph, which was shown to outperform other popular graph-based short read genotyping approaches (Lemay et al. 2022). Following filtering, which discarded variants with excessive genotype missing rate (>5%), heterozygosity (>5%) and low minor allele frequency (<5%), 39,546 SVs genotyped from short reads were available for eQTL analysis (Fig 1B). In parallel, Illumina short read data was used for SNP discovery, resulting in 2,396,948 SNPs included in the eQTL analysis.

### A pangenome reference improves transcript expression quantification for eQTL analysis

One of the key steps in eQTL analysis is the accurate quantification of gene expression. Sequence variation between reference genomes and the actual genotypes used for the generation of expression data can lead to quantification errors, a phenomenon often referred to as ‘reference sequence bias’ (Sibbesen et al. 2023).

To test the effect of potential bias on gene expression quantification in *Brassica napus* and its impact on eQTL analysis, we compared transcript abundance derived from a linear reference based (Kallisto) and a pangenome graph based (RPVG) approach. The pangenome graph reference was constructed using SVs identified and genotyped from long read data combined with SNPs called from short reads. For each transcript, we calculated the correlation between read counts estimated by the two methods across 50 samples (Fig 2A). We then extracted transcripts with Pearson correlation below 0.75 and an equal number of transcripts with highest correlation coefficients. If genomic variation had an appreciable effect on expression quantification, we would expect the transcripts with low measurement concordance across methods to be over-represented in variants. Indeed, we observed a statistically significant enrichment with a permutation test (Fig 2B and Fig 2C) of variants in transcripts with correlation below 0.75. Conversely, the highly correlated transcripts were depleted in variants (Fig S1 and Fig S2). A very similar result was obtained when we used transcripts per million (TPM) instead of counts as a measure of expression. We then tested if the results of the eQTL analysis were also affected. To that end, we performed eQTL detection twice, using the same 50 samples and a similar procedure, with the only difference being the expression quantification method (linear versus graph-based, Fig S3). We found that eQTL transcripts identified only in Kallisto-quantification based analysis were over-represented in variants (Fig S4) compared to those identified only from RPVG-based analysis and those found in both. We concluded that using a pangenome graph reference likely leads to better quantification, therefore RPVG-based expression was selected for the subsequent analysis.

**Figure 2.**
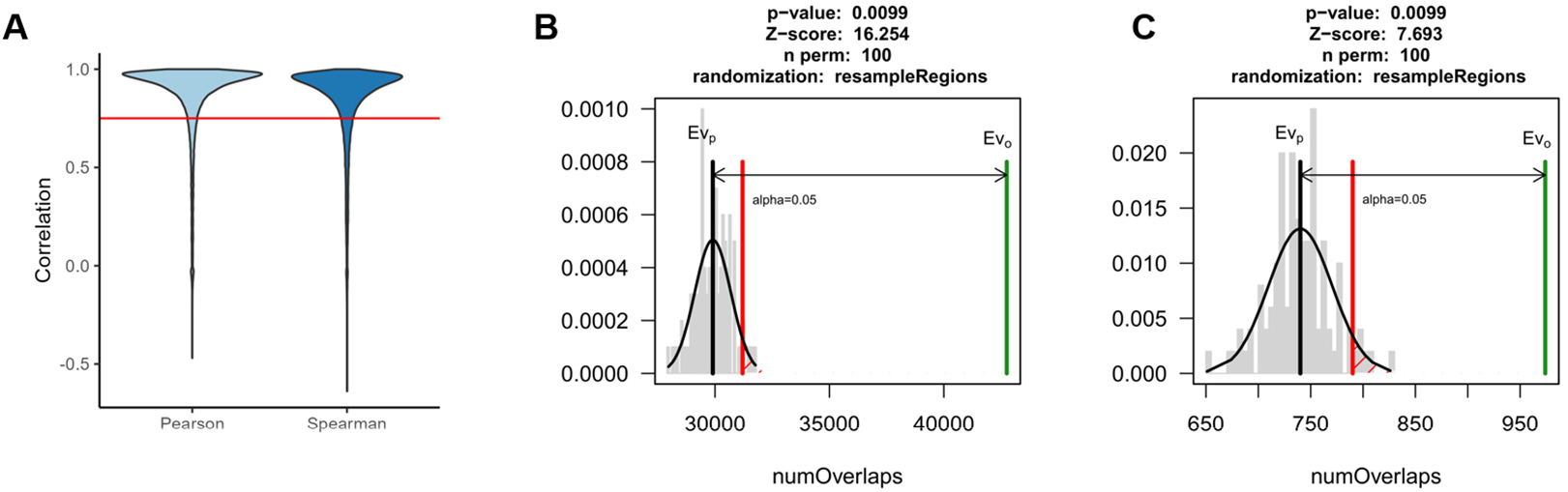
Comparison of linear reference and graph transcript expression quantification approaches. Transcripts with low concordance between Kallisto and RPVG results and overrepresented in genomic variants. **(A)** Pearson and Spearman correlation between Kallisto and RPVG quantification across 50 samples. Red line – 0.75 cutoff used to define transcripts tested for over-representation of variants. Permutation test results: **(B)** transcripts with correlation coefficient below 0.75 are significantly overrepresented in SNPs and **(C)** transcripts with correlation coefficient below 0.75 are significantly overrepresented in SVs. Green line – observed value, grey line - mean of permutation results, red line – significance threshold.

### Gene-proximal structural variants can be linked to gene expression regulation

Final eQTL detection was performed using SNPs, SVs genotyped from short reads and gene expression data representing young leaves at 5-6 leaf stage from 100 homozygous inbred lines. We focused the analysis on lead (identified by lowest p-value) eQTL variants found in proximity of their target genes (cis-eQTLs). Due to the high density of genes in the *B. napus* genome (mean distance between adjacent genes is ∼3,500 bp) we defined cis-eQTL variants as variants located in/overlapping promoter (3,000 base pairs (bp) upstream from the transcription start site (TSS)), exons, introns or regions immediately downstream (3,000 bp downstream from the transcription termination site (TTS)) of their target genes. Using these criteria we identified 240 SV- and 5,668 SNP-eQTLs (Table S1 and Table S2). The proportion of SVs among eQTL variants was higher (4.2%) than among all variants used for the analysis (1.6%). For 36% of SV-eQTL transcripts, no significant associations between any SNP and the transcript were detected, suggesting that these eQTL-SVs are not in linkage disequilibrium with SNPs. For 19% of SV-eQTL transcripts at least one SNP had an equal lowest p-value to the SV (lead SNP and lead SV; in such cases eQTL was retained in both SNP-eQTL and SV-eQTL sets). In the text below we refer to SNP-/SV-eQTLs when discussing variant-transcript pairs and eQTL-SNPs/SVs when referring to variants only.

*Brassica napus* is an allotetraploid, which arose upon the hybridisation of *Brassica rapa* and *Brassica oleracea*. It displays diploid-like behaviour during meiosis and the A and C subgenomes are sufficiently diverged, making it amenable to analysis using bioinformatic methods developed for diploids. However, the asymmetry between subgenomes also opens some interesting questions about their individual contributions to trait regulation. For example, in a recent report, more SNP-eQTLs were identified on the A subgenome compared to the C-subgenome (Tan et al. 2022). Our results agree with this finding. However, we also observed a higher proportion of SV-eQTLs on the C subgenome (Fig 3A, Chi-squared test < 0.001), in line with its higher TE content (Chalhoub et al. 2014) and possibly suggesting asymmetric contributions of SNPs and SVs to cis-regulation in the subgenomes of *B. napus*.

**Figure 3.**
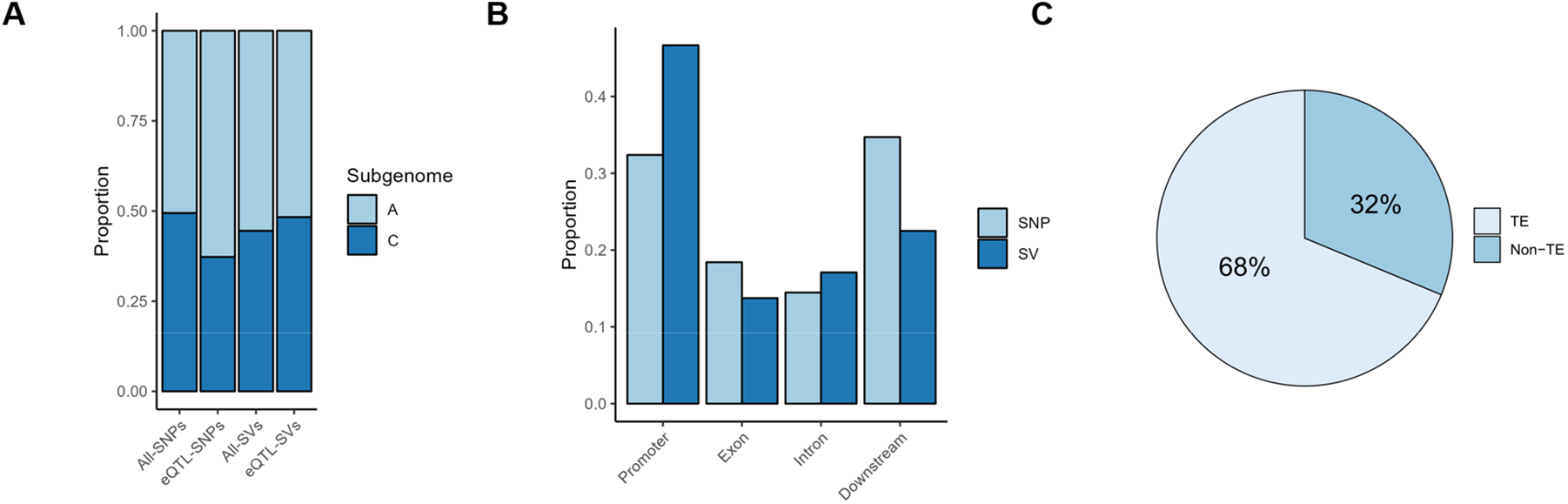
Distribution of eQTLs relative to genomic features. **(A)** eQTL-SNPs and eQTL-SVs are differently distributed across the two subgenomes of *Brassica napus*. **(B)** A higher proportion of eQTL-SVs is found in promoter regions. **(C)** Almost 70% of eQTL-SVs variants have similarity to TEs.

### Majority of cis-eQTL-SVs have similarity to transposable elements

We investigated the distributions of eQTL SNPs and SVs in relation to transcript feature locations. Compared to SNPs, a higher proportion of eQTL-SV were found in promoters (Fig 3B). Overall, a high relative prevalence of SVs upstream of the TSS was previously observed and linked to Class II (DNA) transposable element activity, which can perhaps be explained by easier accessibility of these regions (Fuentes et al. 2019; Han et al. 2013). Indeed, we observed that 68% of eQTL-SVs have similarity to DNA transposable elements (Fig 3C, Fig 4A).

**Figure 4.**
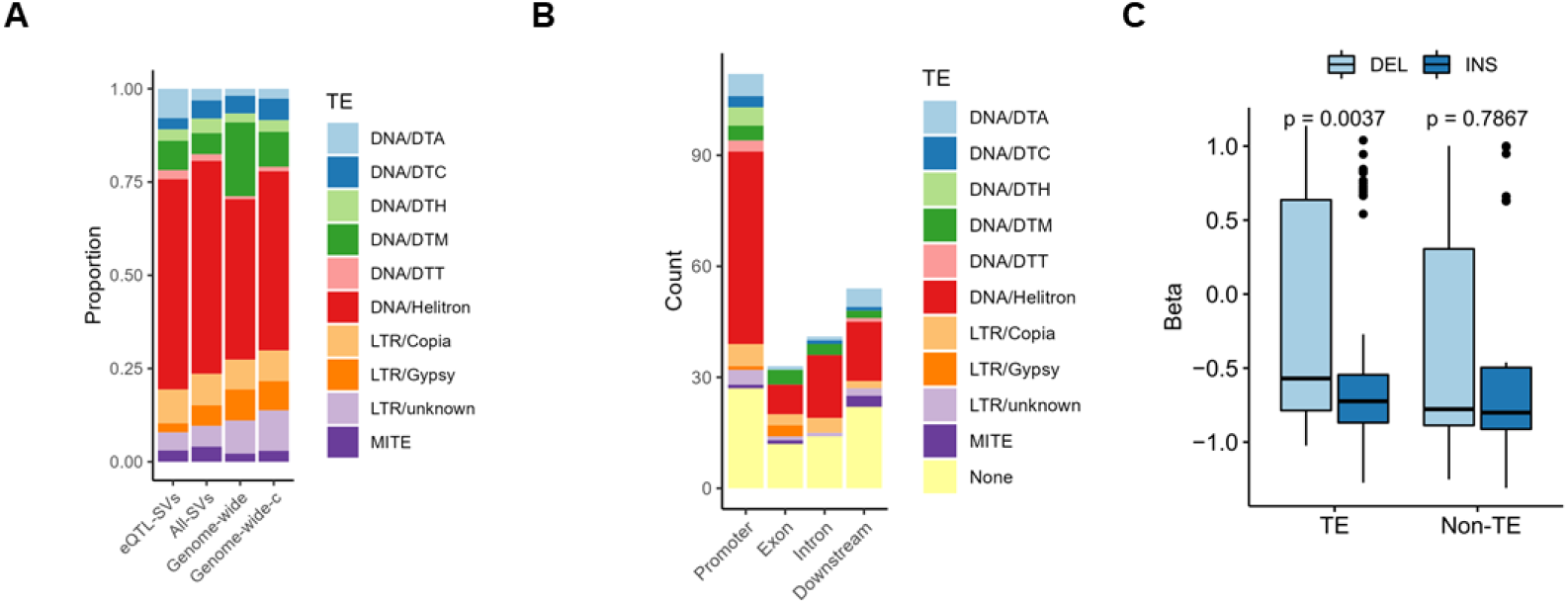
Similarity of SVs to known transposable elements. **(A)** A high proportion of TE-related eQTL-SVs has similarity to Helitrons compared to Genome-wide (based on counts of TEs annotated by GenomeMasker) and Genome-wide-c (based on counts of TEs annotated by GenomeMasker after merging overlapping elements of the same family). **(B)** A high number of promoter-associated eQTL-SVs has similarity to Helitrons compared to eQTL-SVs related to other genomic features. **(C)** TE insertions are associated with decreased expression compared to deletions.

We found that 57% of eQTL-SVs have similarity to Class II (DNA) transposons, 11% to Class I (RNA) transposons and 32% had no detectable similarity to TEs identified in the *B. napus* genome. Among the TE-related eQTL-SVs, the most common TE family was Helitron and the proportion of Helitrons among eQTL-SVs was higher than observed genome wide (Fig 4A). Helitrons were also the most abundant class of TEs found in promoter-located eQTL-SVs (Fig 4B). Overall, TE insertions had a greater negative impact on gene expression than deletions (Fig 4C, Fig S5). Together these results suggest that transposable elements, especially Helitrons, contribute to gene expression diversity in *B. napus*.

Previous eQTL studies have reported a relationship between effect size (Beta) and allele frequency, with SVs associated with higher effect sizes found at lower frequencies in the population (Castanera et al. 2023; Uzunović et al. 2019). We observed a similar pattern in our data for both eQTL-SVs (Fig 5, Figs S6-S9) and SNPs (Fig S10). These results are in line with the expected deleterious effects of rare alleles (Lye et al. 2022) and further support the high quality of our variant and eQTL calls. We observed no significant difference in Beta (Fig S11) and variance (Fig S12) explained by lead eQTL-SNPs and SVs.

**Figure 5.**
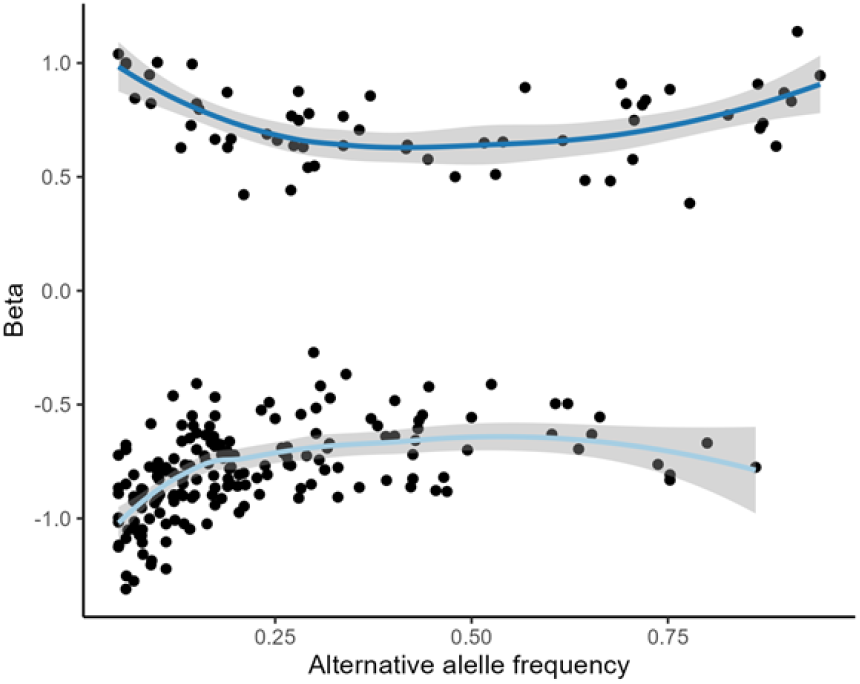
Relationship between alternative allele frequency and effect size (Beta). SVs with higher effect size have a lower frequency in the population.

Out of 233 SV-eQTL transcripts identified, 93% had homologues in the Arabidopsis genome. Gene ontology enrichment analysis did not indicate an over-representation in specific processes or functions. However, some transcripts were annotated with functions related to important traits, including stress response (Fig 6A), morphogenesis (Fig 6B) and flowering. The results suggest that SV-driven gene expression variation could contribute to the phenotypic diversity observed in the field.

**Figure 6.**
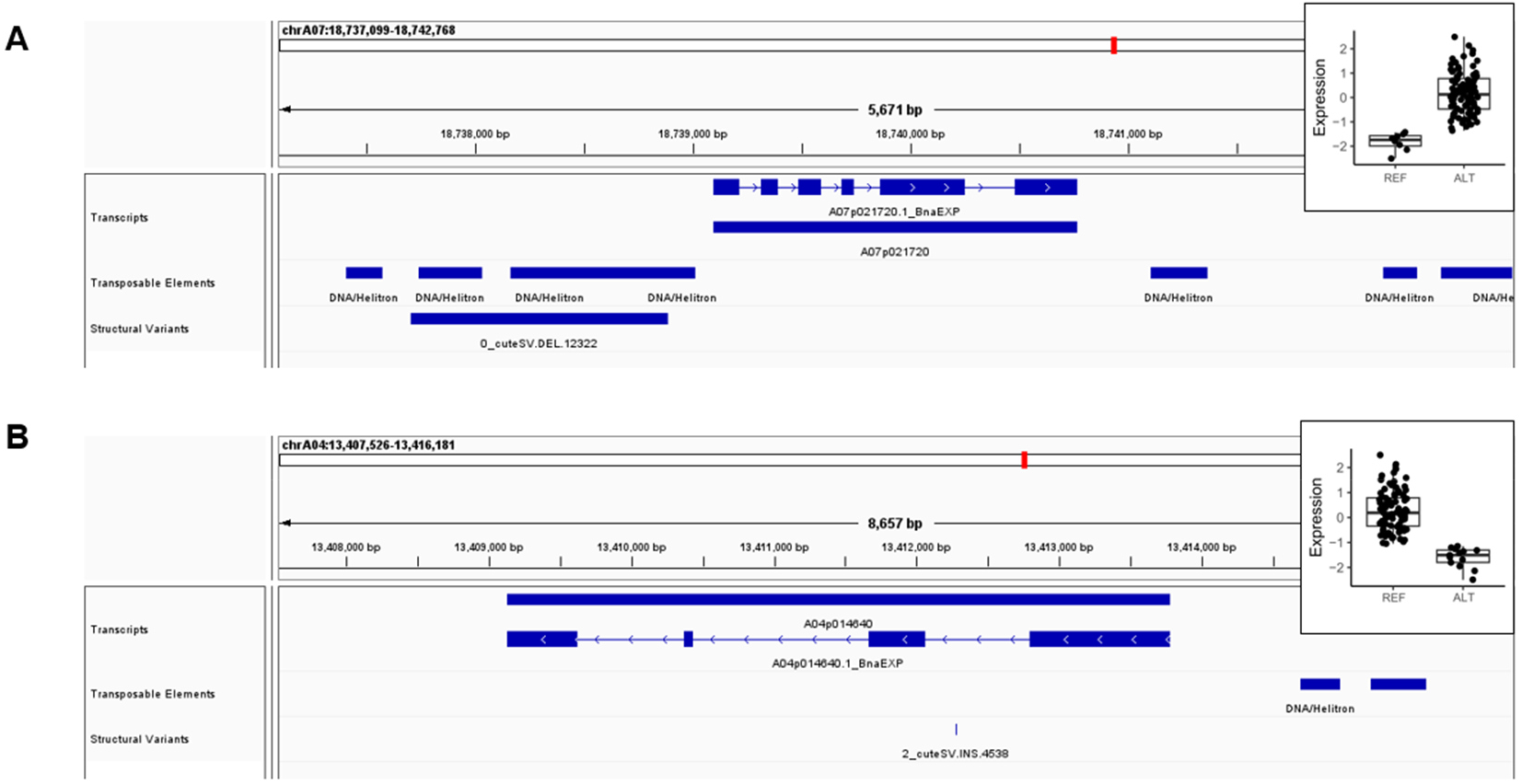
Example of SVs associated with different gene expression levels. **(A)** Deletion of Helitron TE in promoter region is associated with an increase in gene expression. Arabidopsis homologue (SNRK2.8) is involved in response to osmotic stress. **(B)** Insertion of Helitron TE in the first intron is associated with decreased expression. Arabidopsis homologue (SAW2) is involved in leaf morphogenesis. Expression is reported after inverse normal transformation. REF= reference allele, ALT= alternate allele.

## Discussion

We used a pangenome graph approach to discover SNPs and SVs associated with differences in gene expression in young leaves of winter oilseed rape. The pangenome graph was used for structural variant genotyping, but also gene expression quantification.

Our results suggest that bias in transcript expression quantification resulting from a single reference approach can lead to false eQTL discovery, which can be alleviated by the use of a pangenome graph reference. Interestingly, in parallel work we show that genotyping errors may be less of a concern; SVs discovered from long reads mostly either genotype correctly from short reads or fail to genotype altogether (Yildiz et al. unpublished data). However, a failure to genotype does reduce the pool of SVs available for association studies, as exemplified here, with less than half of the initially discovered SVs being successfully genotyped from short reads and included in the downstream eQTL analysis.

We found that the majority of identified *B. napus* eQTL-SVs sequences have similarity to transposable elements. The finding is in line with reports in other crops, including rice and *B. rapa*, where transposable element insertion polymorphisms (TIPs) were shown to contribute to phenotypic and gene expression variation (Cai et al. 2022; Castanera et al. 2023).

Compared to eQTL-SNPs, eQTL-SVs are more likely to be found in promoter regions of genes. The preference of certain transposable elements for insertion into open chromatin regions and especially promoters could make them particularity suited for the rewiring of regulatory networks (Barro-Trastoy and Köhler 2024; Cao et al. 2023; Fuentes et al. 2019). In *B. napus*, the highest number of eQTL-SVs had sequence similarity to Helitrons, likely reflecting their overall high abundance in the genome. Interestingly, Helitrons have also been shown to play important roles in modifying gene regulation in genes involved in endosperm development and response to herbivory (Barro-Trastoy and Köhler 2024).

Overall, TE insertions had a more negative impact on gene expression compared to TE deletions. This pattern was not observed for non-TE SVs. The presence of TEs is known to be associated with increased DNA methylation, which can have a silencing effect on gene expression (Hollister and Gaut 2009). The stronger negative effect of TE insertions suggests that, at least to some extent, epigenetic silencing mechanisms may be involved.

Finally, functional annotation of SV-eQTL transcripts suggests the involvement of some SVs in modulating important biological processes such as stress responses, flowering and morphogenesis. Due to the highly duplicated nature of the *B. napus* genome, owing to whole genome triplication in the ancestral species of Brassica and a more recent allopolyploidization (Cheng et al. 2014), predicting the impact of SVs on traits is not straightforward, even when associated with gene expression differences. Encouraging examples nevertheless exist. For example, despite the presence of multiple homologues, a deletion within the second intron of a *B. napus* FLOWERING LOCUS T homologue was associated with altered flowering time (Vollrath et al. 2021).

Our study highlights the contribution of structural variations to gene expression regulation and the utility of pangenome graph for eQTL analyses. Even using a moderate sample size (n=100), we identified an appreciable number of SVs associated with differences in gene expression. Expanding the sample size and including additional organs and developmental stages will likely result in the identification of many more SVs affecting gene regulation.

## Methods

### Material selection

A total of 100 genetically diverse, elite inbred winter oilseed rape breeding lines from the commercial breeding programme of Norddeutsche Pflanzenzucht HG Lembke (NPZ KG, Hohenlieth, Germany) were used in the study. All 100 lines were used for short-read sequencing. Based on genetic diversity analysis using genome-wide SNPs called from the short-read data, a subset of 57 lines representing the total genetic diversity of the full collection was selected for long-read sequencing. Single plants from each inbred line were harvested for the short and long-read sequencing, respectively.

### Short read genomic and mRNA-Seq sample preparation and sequencing

Plants were grown in a climate-controlled growth chamber with 16 hour day (16°C) and 8 hour night (12°C) cycles. Leaf samples were harvested simultaneously for all genotypes after 30 days at the 5-6 leaf stage, immediately flash-frozen in liquid nitrogen and stored at −80 °C until DNA/RNA extraction. Leaf material was then ground to a fine powder in liquid nitrogen and separated into aliquots for DNA and RNA extraction.

Total genomic DNA was extracted from each sample using the CTAB extraction method of Doyle and Doyle (1990). Total RNA was extracted using the RNeasy Mini Kit (Qiagen, Hilden, Germany) and treated using RNase-free DNAase (Qiagen, Hilden, Germany) to remove DNA. Quantity and quality of RNA samples were checked using a Fragment Analyzer Automated Capillary Electrophoresis system (Advanced Analytical, Heidelberg, Germany). Equimolar RNA/DNA samples were shipped on dry ice to BGI Tech Solutions (Hong Kong) for sequencing.

Whole-genome DNA sequencing was performed with 150nt paired-end reads on the Illumina Hiseq XTen platform. mRNA-Seq was performed on the Illumina HiSeq 4000 platform with 100 nt paired-end sequencing.

### Long read genomic sample preparation and sequencing

Plants were grown in the same conditions as described above, leaves were harvested from plants at the 4-6 leaf stage, flash-frozen and ground to a fine powder using a mortar and pestle. High-molecular-weight DNA was isolated and sequenced using a modified protocol from Chawla et al (2021). Briefly, 11 mL of pre-heated lysis buffer (1% w/v PVP40, 1% w/v PVP10, 500 mM NaCl, 100 mM TRIS pH8, 50 mM EDTA, 1.25% w/v SDS, 1% (w/v) Na_2_S_2_O_5_, 5 mM C_4_H_10_O_2_S_2_, 1% v/v Triton X-100) were added to 1.2-1.5 g of tissue and incubated for 30 min at 37°C in a rotator. 11 μl of RnaseCocktail (ThermoFisher, ref AM2288) were added and the lysate was incubated in a rotator at 37°C for 20 minutes. 110 μl of ProteinaseK (ThermoFisher, ref QS0511) were then added and samples were incubated in a rotator at 37°C for a further 20 minutes. 4 mL of 5 M potassium acetate were added to the lysate after cooling, then mixed by inversion 20 times, incubated for 10 min on ice and pelleted by centrifugation at 4°C, 4250 RCF for 10 min. Finally, magnetic beads were used to recover the HMW-DNA, washed twice with 70% ethanol and incubated with TE buffer for 10 minutes at 37°C to release the DNA from the beads into the buffer. 1 to 3 μg of DNA were used for library preparation with the ligation sequencing kit SQK-LSK109, according to manufacturer’s recommendations, and loaded onto an Oxford Nanopore MinION flow cell for sequencing.

### SV identification, genotyping and quality control

SV discovery was performed using a previously established protocol (Yildiz et al. 2023). In short, reads were mapped to the Express 617 v1 (Lee et al. 2020) reference genome using minimap2 (Li 2018) v2.24-r1122, structural variants (SV) for each of the 57 lines were called using cuteSV (Jiang et al. 2020) v1.0.13. SVs were merged using Jasmine (Kirsche et al. 2023) v1.1.5 and re-genotyped using cuteSV. Variants were filtered to retain insertions and deletions with genotype missing call rate below 5% and heterozygous genotype call rate below 5%. Insertions and deletions constitute a vast majority of SVs identified between *B. napus* genotypes (Yildiz et al. 2023) and are associated with lowest call error rates among all variant types (Jiang et al. 2021). SVs with excessive heterozygous calls were removed, as they were previously shown to be caused by genotyping errors in samples, which were expected to be highly homozygous, as is the case for *B. napus* inbred lines (Yildiz et al. 2023). Finally, SVs longer than 20,000 bp were also removed, as we showed previously that they constitute only a very small percentage of SVs in *B. napus* (Yildiz et al. 2023) and the main focus of this study was on smaller insertions and deletions and their impact on cis-regulation.

### SNP identification

Reads were mapped to the Express 617 v1 reference genome using BWA-MEM2 (Vasimuddin et al. 2019) v2.2.1. SNPs were identified using bcftools (Danecek et al. 2021) v1.15.1 (multiallelic-caller) with minimum read mapping quality of 10. Similarly to the SVs, variants with genotype missing call rate below 5%, heterozygous genotype call rate below 5% and minor allele frequency above 5% were retained.

### Gene expression quantification using RPVG and Kallisto

A pangenome graph was built using vg (Garrison et al. 2018) v1.4.30 autoindex, based on the Express617 v1 reference genome sequence and using SNPs and SVs which passed the quality control filtering steps described above. mRNA-Seq reads were mapped to the graph using vg mpmap. The mappings were passed to RPVG for quantification. For each sample, RPVG outputs quantification results along with haplotype probabilities above a certain threshold. The per-sample results were filtered to retain only haplotypes with the highest probability for each gene. Further, only genes for which the haplotype could be assigned in all samples were retained. Transcripts per million (TPM) values were extracted directly from the RPVG output. Kallisto v0.44.0 was used for quantification using transcripts extracted from the Express617 v1 assembly. TPM values were extracted directly from Kallisto outputs. Transcripts quantified with Kallisto, which could not be assigned a haplotype by RPVG for all samples, were removed prior to comparisons.

### Comparison of quantifications using RPVG and Kallisto

Pearson and Spearman correlations were calculated for each gene across 50 samples. Transcripts with Pearson correlation below 0.75 were tested for over-representation of SNPs and SVs using a permutation test implemented in regioneR (Gel et al. 2016) v1.26.1. All transcripts quantified by RPVG in all samples were used as a universe for resampling in 100 iterations.

To test whether quantification bias can affect eQTL identification, the same eQTL analysis pipeline (as described below) was used, with the only difference being the method of transcript expression quantification. We then extracted transcripts that (a) were only identified in the analysis performed using Kallisto, (b) were only identified in the analysis performed using RPVG, and (c) were identified in both analyses. These were tested for the over-representation of variants using a permutation test implemented in regioneR. All eQTL transcripts identified in both analyses were used as the universe for resampling.

### eQTL identification

Only transcripts with a mean TPM value of at least 0.1 and expression of at least 1 TPM in at least two samples were retained. The expression matrix was quantile normalized and transformed using inverse normal transformation. Five top principal components (PCs) identified from SNP data and top components identified from expression data using the Elbow method of PCAforQTL (Zhou et al. 2022) v0.1.0 were used as covariates in matrixEQTL (Shabalin 2012) v2.3. eQTL analysis was performed jointly for SNPs and SVs, with FDR adjusted p-value < 1e-6. Prior to eQTL analysis, variants were further filtered to remove variants with minor allele frequency below 5%. Lead variants were identified by lowest p-value. When SNPs and SVs had equal lowest p-value both were retained.

### Transposable element annotation

Transposable element library for the Express 617 genome assembly was generated using EDTA (Ou et al. 2019) v2.0.1. Transposable elements were annotated using RepeatMasker v4.1.2. Sequences of insertions and deletions were extracted and compared against the TE library using BLASTn (Camacho et al. 2009) v2.13.0 (e-value cut-off 1e-5). Top BLAST matches were used to assign SV sequences to TE families.

### Arabidopsis homologue identification and GO term annotation

Homologue identification was performed using a previously developed method (Golicz et al. 2021). Briefly, protein sequences of *B. napus* transcripts were compared against *Arabidopsis thaliana* proteome database using BLASTp v2.13.0 (e-value cut-off 1e-5). Best *A. thaliana* matches of *B. napus* transcripts were identified as homologues. GO annotation was performed by transferring TAIR GO annotation of *A. thaliana* to the *B. napus* homologous genes.

## Data access

All raw data generated in this study have been submitted to the NCBI BioProject database (https://www.ncbi.nlm.nih.gov/bioproject/) under accession number PRJNA1086556.

## Competing interest statement

The authors declare no competing interests.

## Acknowledgments

This work was supported by the Alexander von Humboldt Foundation in the framework of Sofja Kovalevskaja Award to AAG, grant 031B0888G from BMBF to AA, RJS and by the BMBF-funded de.NBI Cloud within the German Network for Bioinformatics Infrastructure (de.NBI) (031A532B, 031A533A, 031A533B, 031A534A, 031A535A, 031A537A, 031A537B, 031A537C, 031A537D, 031A538A). This work was performed with support from Justus Liebig University Bioinformatics Core Facility (BCF). We thank Regina Illgner and Stavros Tzigos for the technical support and Dr. Paul Knight for critical reading and feedback during the manuscript preparation.

## Author Contributions

Conceptualization: AAG; Methodology: AAG, SFZ; Formal analysis: AAG, GY, SFZ; Investigation: AAG, SW, SFZ; Writing - Original draft: AAG, SFZ; Writing – Review and editing: AAG, GY, SW, TK, AA, RJS, SFZ; Visualization: AAG; Supervision; AAG; Project administration: AAG, RJS, SFZ; Funding Acquisition: AAG, AA, RJS.

## Notes

### Competing Interest Statement

The authors have declared no competing interest.

